# A BubR1-independent pathway for CENP-E targeting to the outer corona of kinetochores

**DOI:** 10.1101/2023.05.10.540161

**Authors:** Thibault Legal, Calum Paterson, Agata Gluszek, Owen R. Davies, Julie P.I. Welburn

## Abstract

For chromosome segregation to take place, unattached kinetochores expand in early mitosis, forming a fibrous structure called the fibrous corona that is captured by microtubules. The corona is assembled from the RZZ complex, Spindly, CENP-E and the Mad1/Mad2 spindle assembly checkpoint proteins. CENP-E aligns chromosomes along the mitotic spindle by moving them to the plus ends of microtubules. Here, we show that CENP-E is recruited to the outer corona independently of BubR1 in a dynein-dependent fashion. We determine the structure of this domain and show that a conserved loop is essential for CENP-E targeting to the outer corona. We show that both domains are essential for CENP-E recruitment to unattached kinetochores. We also report that the kinetochore-targeting domain of CENP-E contributes to the recruitment of the RZZ complex, Mad1 and Spindly, providing a feedback loop to assemble the outer corona. In this study, we propose that CENP-E uses 2 pathways to target to the kinetochore, which allows it to optimize kinetochore capture by microtubules for chromosome alignment and mitotic progression.

---

Chromosome alignment and biorientation is essential to ensure faithful portioning of sister chromatids to daughter cells. The kinetochore of chromosomes captures microtubules emanating from opposites poles of the spindle to facilitate alignment and biorientation. Once the spindle checkpoint is satisfied, chromosome segregation takes place. The kinetochore is a key multiprotein structural assembly that connects the centromere of chromosomes to dynamic microtubules (Cheeseman and Desai, 2008). The kinetochore consists of an inner constitutive CCAN complex, connecting to CENP-A and CENP-C which specify the centromeric chromatin, and the KMN network at the outer kinetochore, which connects to incoming microtubules (Musacchio and Desai, 2017). Some proteins are dynamically localized at the kinetochore as its attachment state to microtubules changes (Hoffman et al., 2001). In prometaphase, the spindle assembly checkpoint proteins associate with the outer kinetochore and sense the attachment state of kinetochores to inhibit the spindle checkpoint until biorientation occurs. The fibrous corona is a low-density structure recruited outside of outer kinetochores in the absence of microtubules in early mitosis (Jokelainen, 1967; McEwen et al., 1998; Rieder, 1982). The fibrous corona is assembled from the oligomerization of the RZZ complex (ROD, Zwilch and ZW10) and is dependent on the dynein adaptor Spindly and the kinase Mps1 activity (Pereira et al., 2018; Rodriguez-Rodriguez et al., 2018; Sacristan et al., 2018). CENP-E is present at the outer corona, along with Mad1 and Mad2. Proteomic studies also indicate CENP-E is in complex with RZZ/Spindly/Mad1 cluster (Samejima et al., 2015). The fibrous corona expands to maximize the attachment of kinetochores to microtubules and then disassembles upon kinetochore-microtubule attachment and biorientation (Hoffman et al., 2001; Magidson et al., 2015). Removal of the corona and checkpoint silencing are coupled and depend on dynein removal of Spindly-RZZ Mad1/Mad2 and CENP-E (Howell et al., 2001).

The CENP-E motor (kinesin-7) plays a critical role in aligning polar chromosomes that failed to align using other pathways (Craske and Welburn, 2020; Maiato et al., 2017). Inhibition or depletion of CENP-E results in chromosomes accumulating at the spindle poles and unable to congress, leading to aneuploidy (Wood et al., 1997; Bennett et al., 2015). The CENP-E localizes to the outer kinetochore and the fibrous corona, and remains at kinetochores after fibrous corona disassembly throughout anaphase (Cooke et al., 1997). The kinetochore-targeting region of CENP-E resides in its C-terminal region (Chan et al., 1998). Recent work revealed this C-terminal region of CENP-E binds to the pseudokinase domain of the checkpoint protein BubR1 (Ciossani et al., 2018; Legal et al., 2020). The recruitment of CENP-E to kinetochores via BubR1 upon activation of the spindle checkpoint is rapid-within minutes. The BubR1-dependent recruitment of CENP-E is essential for correct chromosome alignment and segregation. However, CENP-E is also recruited via second pathway, independently of BubR1 to unattached kinetochores. This recruitment is slower (Legal et al., 2020), presumably the time for assembly of the outer corona being rate-limiting. However, the molecular basis for CENP-E targeting to the outer corona and its function there remain unclear. Here we determine the structure of the corona-targeting domain of CENP-E and define the molecular requirements and dependency for its targeting. We show that both the BubR1 and corona-targeting domains of CENP-E contribute to the recruitment of the RZZ complex, Spindly and Mad1, and that disassembly of the complex is key to complete biorientation.

## CENP-E recruitment to and removal from kinetochores depends on Dynein

The RZZ complex and Spindly self-assemble into this fibrous corona and recruit CENP-E and Mad1. Dynein is also present at unattached kinetochores (Raaijmakers et al., 2013). To further characterise the BuBR1-independent recruitment pathway to kinetochores (Legal et al., 2020), we arrested the cells in nocodazole for 2 hours and depleted either BubR1, dynein or both (Fig S1A). Dynein heavy chain depletion does not affect the integrity of the outer corona. Depletion of BubR1 did not significantly decrease CENP-E at kinetochores, in agreement with our previous work. Depletion of Dynein led to a modest reduction in CENP-E at kinetochores (Fig 1A, B). Depletion of both BuBR1 and Dynein led to a strong reduction in CENP-E recruitment to unattached kinetochores, with very little CENP-E remaining at kinetochores (Fig 1A, B). In the absence of Dynein, CENP-E at kinetochores was also reduced in both prometaphase and metaphase cells, indicating Dynein plays a role in CENP-E recruitment to kinetochores (Fig 1C, D). To further test whether dynein and CENP-E work together, we inhibited endogenous CENP-E in its rigor state using the allosteric inhibitor GSK923295 for 30 minutes. Thus CENP-E remains bound to microtubules and is unable to take any steps. CENP-E was absent from kinetochores and accumulated to the spindle poles, marked by centrin. However, when Dynein was depleted by siRNA, CENP-E did not accumulate at the poles. The levels of CENP-E at kinetochores were strongly reduced and some residual CENP-E was observed on the spindle close to the spindle poles marked by centrin, possibly due to CENP-E in rigor state binding to microtubules (Fig 1E, F). Overall, this indicates that while Dynein removes CENP-E once biorientation has occurred (Howell et al., 2001; Wojcik et al., 2001), CENP-E also depends on dynein for its targeting to the fibrous corona.

**Figure 1:**
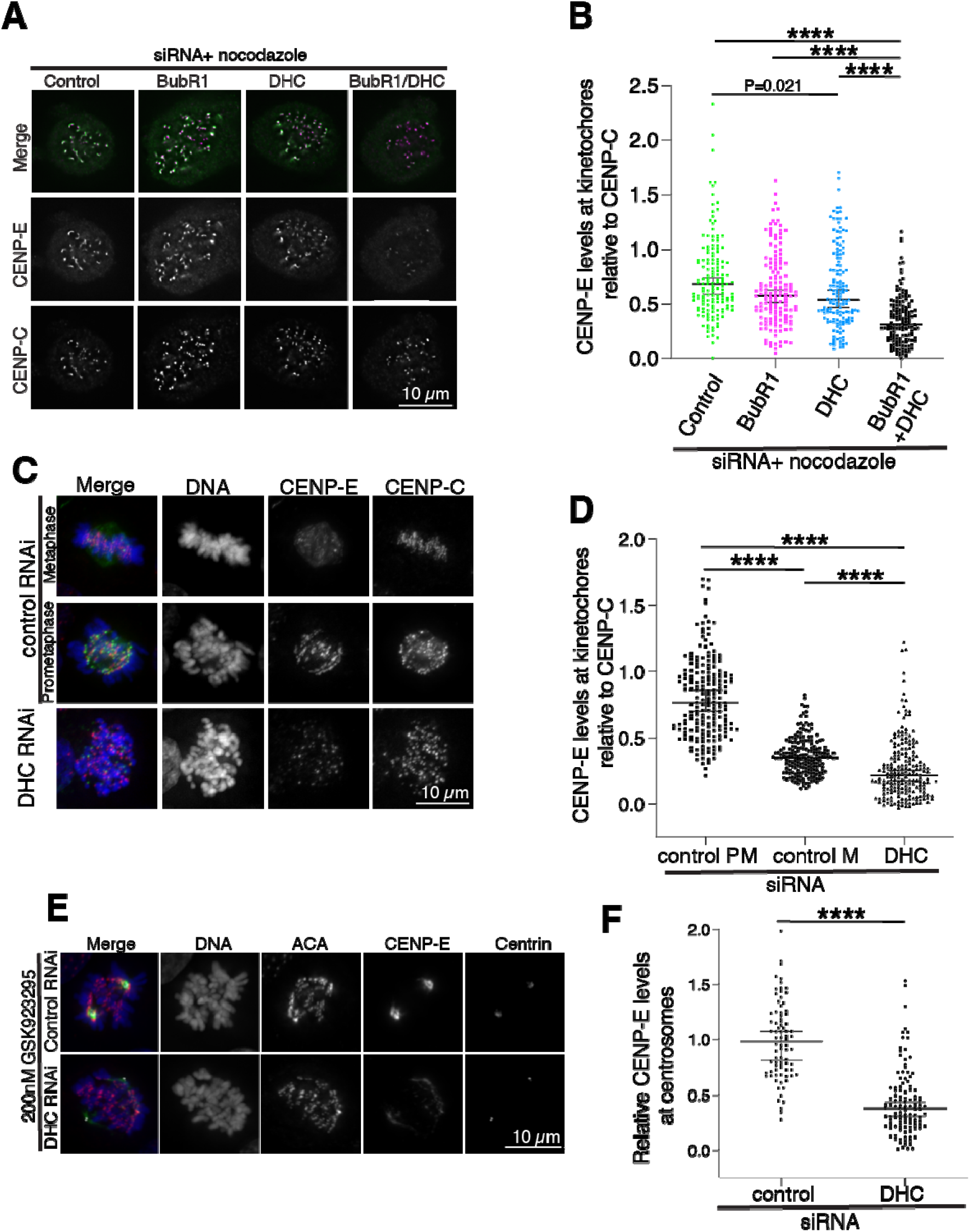
CENP-E recruitment to unattached kinetochores depends on two distinct pathways involving BubR1 and dynein. (A) Representative immunofluorescence images of HeLa cells transfected with CENP-E constructs after depletion of BubR1, Dynein Heavy chain (DHC) or BubR1/DHC and treated with nocodazole for 2 hours. The cells were stained for endogenous CENP-E, CENP-C and DNA. (B) Scatter dot plot showing quantification of CENP-E intensity normalized to CENP-C, in the absence of endogenous CENP-E. Median and 95% confidence interval are presented. Cells were treated for 2 hours with nocodazole. Each point represents the intensity of CENP-E over CENP-C at one kinetochore. Black line represents the mean and whiskers represent the standard deviation. Asterisks indicate ordinary Kruskal-Wallis test significance value. ****P<0.0001, *P=0.02. n=150 kinetochores for each condition. (C) (D) Scatter dot plot showing quantification of CENP-E intensity normalized to CENP-C for cells in prometaphase, metaphase or metaphase depleted with Dynein, after treatment with control or DHC siRNA for 48 hours. n=201, 202 and 200 respectively. Median and 95% confidence interval are presented. Asterisks indicate ordinary one-way ANOVA test significance value. ^****^P<0.0001. (E) Representative immunofluorescence images of HeLa cells after depletion with control or Dynein Heavy Chain siRNA and treated with 200nM GSK923295 for 30q minutes. (F) Scatter dot plot showing quantification of CENP-E intensity around spindle poles, median and 95% confidence interval. n= of 83 and 113 cells measured respectively. For each cell, the intensity around both spindle poles marked by Centrin was measured and averaged. Scale bars: 10 μm. Asterisks indicate T-test significance value. ****P<0.0001.

### CENP-E targeting to the outer corona involves a domain distinct from the BubR1-binding domain

Our data indicate that CENP-E is recruited through a pathway separate from BubR1 and dependent on Dynein (Fig 1) (Legal et al., 2020). We thus hypothesized that the corona targeting pathway is distinct from the minimal BubR1-interacting domain which we previously identified as residues 2091-2358. To dissect the minimal requirement for CENP-E targeting to the outer corona, we transfected CENP-E domains fused to GFP in the presence of either nocodazole or the CDK1 inhibitor RO-3306 and nocodazole in a cell line expressing mCherry-CENP-A to mark kinetochores. While in the presence of nocodazole, all outer kinetochore and corona proteins are present, brief treatment with CDK1 inhibitor causes disappearance of the outer kinetochore proteins while the ones associated with the outer corona detach from the kinetochore as a fibrous structure (Sacristan et al., 2018). CENP-E_2091-2358_-GST-GFP, which we previously showed to be recruited to kinetochores through BuBR1, localized to the kinetochore in the presence of nocodazole but not after treatment with the CDK1 inhibitor (Fig 2A). GFP-CENP-E_2055-2450_ targeted to kinetochores but also failed to target to the detachable outer corona under these conditions. However, GFP-CENP-E_2452-2598_ remained present at the outer corona of the outer kinetochore in the presence of the CDK1 inhibitor (Fig 2A). Under these conditions, we then imaged GFP-CENP-E_2452-2598_ alongside the outer corona protein and dynein adaptor Spindly. Indeed GFP-CENP-E_2452-2598_ associates with the detachable corona crescent, co-localizing with Spindly and is separate from the constitutive kinetochore marked by endogenous CENP-C (Fig 2B). Overall, this work reveals that region 2452 to 2598 associates with the fibrous outer corona, where it co-localizes with Spindly (Fig 2B, C).

**Figure 2:**
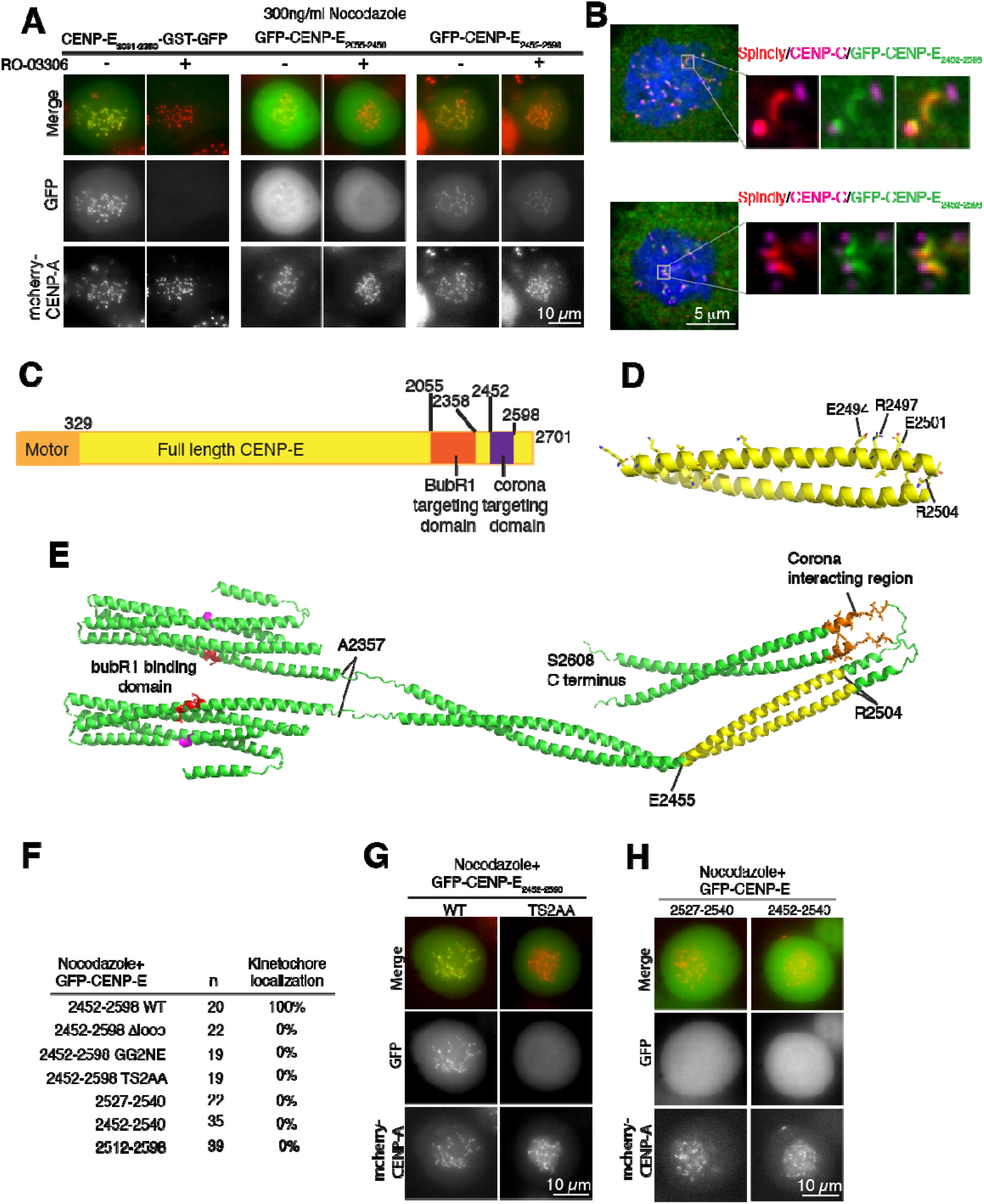
A conserved loop downstream of BubR1 binding domain in CENP-E is essential for targeting to the outer corona. (A) Representative images of live HeLa cells transfected with GFP-CENP-E constructs, treated with 300 nM nocodazole and with or without the CDK1 inhibitor RO-3306, scale bar: 10 μm. (B) Representative immunofluorescence images of HeLa cells transfected with GFP-CENP-E_2452-2598_ and treated with nocodazole and RO-3306, stained with Spindly and CENP-C. Scale bar, 5 μm. (C) Schematic diagram of CENP-E, highlighting the motor (orange) and kinetochore- and corona-targeting domains (purple). (D) X-ray crystallography derived structure of CENP-E_2454-2494_ in cartoon representation. The conserved amino acids are represented in ball and stick mode (PDB:8OWI). (E) Model of the CENP-E_2055-2608_ generated by combining Alphafold2 models of its monomeric kinetochore targeting domain and dimeric outer corona targeting domain. The conserved residues essential for corona-targeting is in stick and ball orange. The domain crystallized (D) is painted yellow. The residues essential for interaction with BubR1 are in red. (F) Quantification of the kinetochore localization of GFP-CENP-E domain mutants related to images in F and G. (G, H). Representative live-cell images of HeLa cells transfected with GFP-CENP-E point (G) or domain (H) mutants and treated with 300 nM nocodazole for 2 hours. Scalebar, 10 μm.

### A flexible loop linker is essential for CENP-E outer corona targeting

Next, we sought to determine the structure of this minimal corona targeting domain. This region contains a highly conserved unstructured sequence, flanked by two predicted α-helices. We expressed and purified CENP-E_2452-2598_ (Fig S1B). We obtained protein crystals that diffracted anisotropically with resolution limits between 2.14 Å and 2.71 Å (Table 1). Structure determination revealed a parallel dimeric coiled-coil in which only the first helical region of the construct was observed in electron density (Fig 2D). The dimeric nature is consistent with the behaviour of CENP-E_2452-2598_ in solution. Crosslinking with BS3 indicates it is a dimer (Fig S1B) and previous work showed this domain was responsible for dimerization of the kinetochore targeting domain of CENP-E_2055-2608_. It is likely that the construct underwent C-terminal degradation during crystallization. The resultant structural model corresponds to amino acids K2454 to S2506 of CENP-E. The structure reveals a tight left-handed coiled-coil, in which the hydrophobic heptad interface includes aromatic stacking of F2478 residues, and with surrounding interactions include salt-bridges between E2470 and K2471 side-chains.

To gain insights into the wider architecture of the corona and kinetochore-targeting domains, we built a molecular model of the CENP-E_2055-2608_ dimer (Fig 2E). In preliminary analyses, Alphafold2 predicted two distinct domains, in keeping with our previous findings (Legal et al., 2020), and thus we built a model of the structure from these two separate regions. As we previously showed that the BubR1-targeting domain is a monomer, we firstly used Alphafold2 to model a region encompassing this domain (amino-acids 2055-2377) as a monomer. This revealed a four-helical bundle structure with high confidence (Fig S2). We then used Alphafold2 to model the subsequent region encompassing the corona-targeting domain (amino-acids 2356-2608) as a dimer, using our newly solved crystal structure as a template. This revealed two consecutive parallel homodimeric coiled-coils (residues 2370-2514 and 2532-2605) that flank the central conserved corona-targeting loop. Part of the last C-terminal coiled coil is also highly conserved (Fig S1C). The structure we determined is part of the coiled coil on the N-terminal side of the conserved loop (highlighted in yellow, Fig 2E). We combined these models to present a model of the complete CENP-E_2055-2608_ dimer. Overall, this model fits with the rotary shadowing of CENP-E_2055-2608_, which revealed the presence of a globular domain adjacent to an extended domain, with overall lengths of 36 nm and 40 nm, respectively (Legal et al., 2020). In the model, the BubR1 binding region is solvent exposed and in close proximity to another highly conserved sequence “DFSE” which may also contribute to BubR1 binding (Fig 2E, magenta).

Given the high conservation of a part of the loop across multiple species in the corona-targeting domain (Fig 2C), we hypothesized that it could be important for recruitment of CENP-E to the outer corona. Mutating T2529 and S2534 to A (TS2AA), or G2532 and G2533 to NE (GG2NE) in CENP-E_2452-2598_ also abolished their recruitment to the corona (Fig 2F, G). We generated a mutant lacking the conserved residues in the loop and the start of the consecutive helix (GFP-CENP-E Δloop), removing amino acids 2527-2538. GFP-CENP-E Δloop showed no localization to the outer corona indicating this part of the loop is critical for its recruitment there (Fig 2F). We then tested if the loop sequence and loop flanked by the neighbouring helices were sufficient for outer corona targeting by transfecting GFP-CENP-E_2527-2540_. The sequences _2527_PLTCGGGSGIVQNT_2540_, the _2452_helix-loop_2540_ and _2512_loop-helix_2598_ were not sufficient for targeting to the outer kinetochore (Fig 2F, H). Overall, while the conserved loop seems essential for CENP-E targeting to the outer corona, it is not sufficient. The flanking helical coiled coils also appear necessary for recruitment to the outer corona.

### CENP-E is targeted to kinetochores through BubR1 and the corona-associated proteins

CENP-E targets to kinetochores through at least 2 pathways (Legal et al., 2020). Thus, we hypothesized that disrupting both the acidic patch which binds to BubR1 using the previously published GFP-CENP-E_2055-2608_ E4A mutant (Legal et al., 2020), and the corona-targeting loop using the TS2AA mutant, should prevent recruitment of CENP-E to unattached kinetochores. Indeed, GFP-CENP-E_2055-2608_ localized to unattached kinetochores in cells treated with nocodazole for 2 hours (Fig 3A, B). The GFP-CENP-E_2055-2608_ E4A and GFP-CENP-E_2055-2608_ TS2AA mutants were still able to target to the outer corona. However, the localization of the double mutant GFP-CENP-E_2055-2608_ E4A TS2A was almost abolished (Fig 3A, B). Next, we tested the recruitment of CENP-E to kinetochores in the context of the full-length protein. Full-length CENP-E wild type and mutants were transfected into HeLa cells and treated with nocodazole to analyze their recruitment to unattached kinetochores (Fig 3C, D). CENP-E-GFP E4A and CENP-E-GFP TS2AA showed a marked reduction in kinetochore localization, while the CENP-E-GFP E4A/TS2AA was not present at the kinetochores. Overall, this indicates there are two parallel pathways that recruit CENP-E to unattached kinetochores.

**Figure 3:**
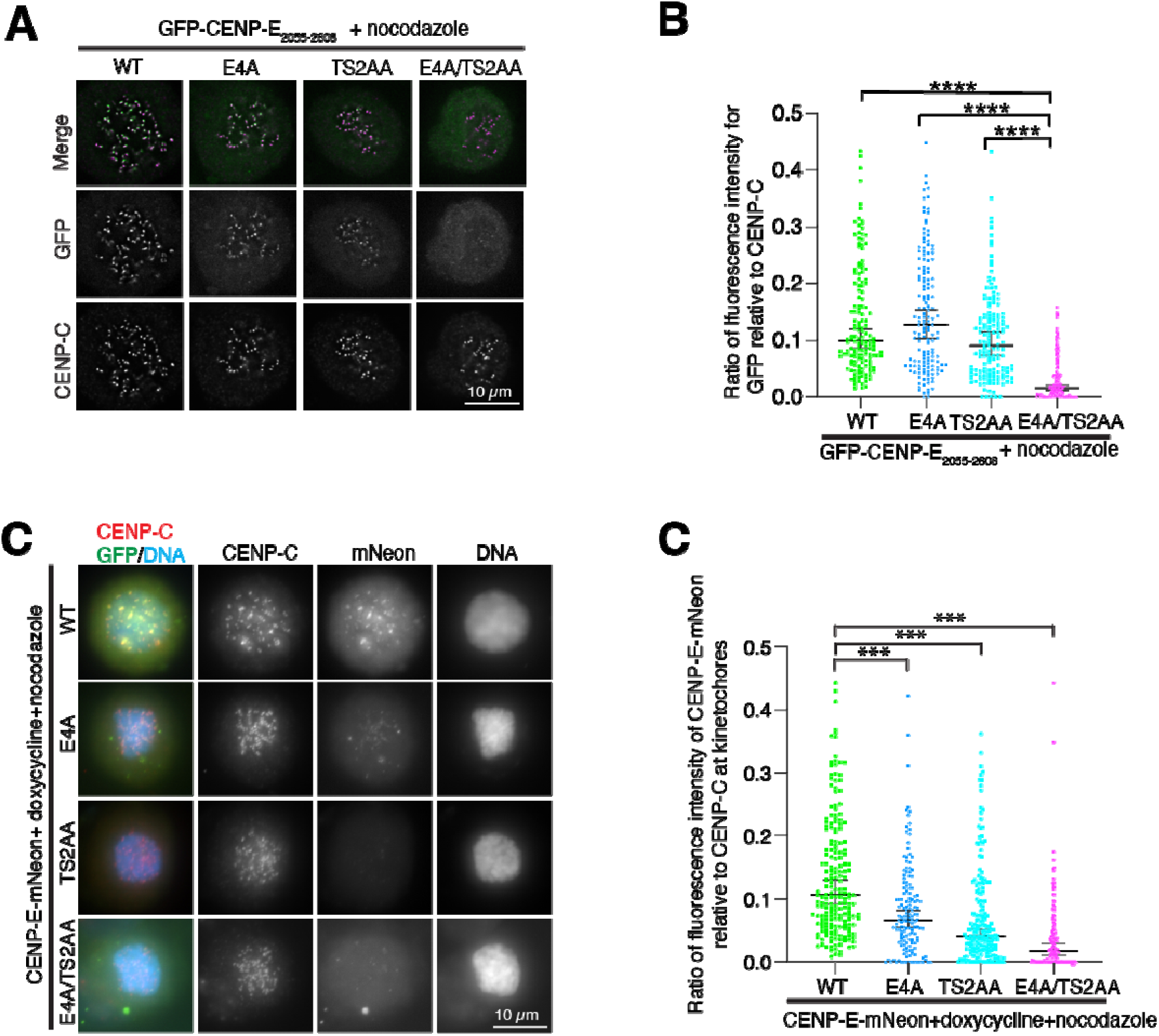
Disruption of the BubR1 and corona interactions motives prevent CENP-E localization to kinetochores. (A) Representative immunofluorescence images of HeLa cells transfected with GFP-CENP-E_2055-2608_ wild type and mutants, treated with 300 nM nocodazole. Scalebar, 10 μm. (B) Scatter dot plot showing quantification of GFP-CENP-E_2055-2608_ intensity relative to CENP-C at kinetochores. The experiment was done twice. Scale bars: 10 μm. n=150 for each condition, bars represent median and 95% confidence interval. **** P<0.0001. Asterisks indicate ordinary Kruskal-Wallis test significance value. (C) Representative immunofluorescence images of HeLa cells transfected with full-length CENP-E-mNeon wild type and mutants, CENP-C and DNA. Scalebar, 10 μm. (D) Scatter plot showing ratio of fluorescence intensity of mNeon relative to CENP-C for CENP-E wild type and mutants at spindle poles (kinetochore numbers for WT n=194, E4A n=148, TS2AA n=197 and E4A TS2AA n=150). Bars represent median and 95% confidence interval. Asterisks indicate ordinary Kruskal-Wallis test significance value, with **** P<0.0001.

### The corona components relocalize with CENP-E_2055-2608_ and inhibit kinetochore biorientation

We previously showed that GFP-CENP-E_2055-2608_ accumulates at the spindle poles and centrioles (Legal et al., 2020). We hypothesized that it may be able to delocalize the other components of the detachable corona and target them to the spindle poles, thereby causing the misaligned chromosomes. Indeed, in the presence of GFP-CENP-E_2055-2608_, Spindly and Mad1 co-localized at the spindle poles and enriched there (Fig. 4A-D). This was also the case for ZW10, a subunit of the RZZ complex that forms the fibrous mesh of the outer corona (unpublished). GFP-CENP-E_2055-2608_ E4A and GFP-CENP-E_2055-2608_ TS2AA still accumulated at the poles, suggesting they make localize there through direct interaction with dynein (Fig. 4E, 1E, F). However, there was a reduction in the levels of the other corona protein components Mad1, Spindly and RZZ, marked by ZW10 (Fig. 4A-D). In the presence of GFP-CENP-E_2055-2608_ E4A TS2AA which impairs both BubR1 and corona binding of CENP-E, Spindly and Mad1 remained associated with kinetochores. They did not associate with GFP-CENP-E_2055-2608_ E4A TS2AA at the spindle poles (Fig. 4). In total, these data indicate that GFP-CENP-E_2055-2608_ interacts with components of the outer corona Mad1, Spindly and the RZZ complex and is able to recruit them directly.

**Figure 4:**
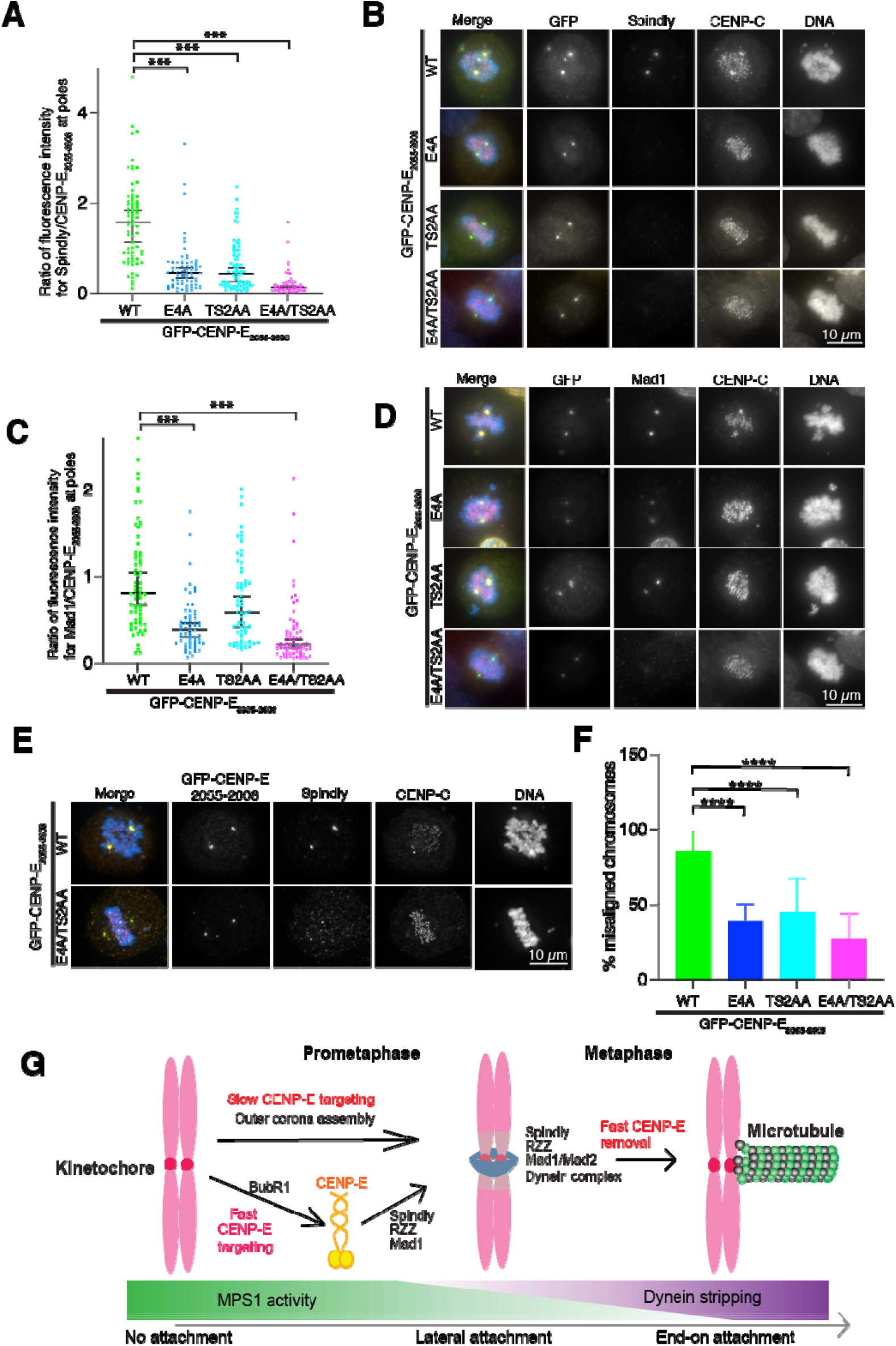
CENP-E_2055-2608_ associates with Spindly, Mad1 and the RZZ complex through the corona binding motif. (A) Representative immunofluorescence images of HeLa cells transfected with GFP-CENP-E_2055-2608_ wild type and mutants, stained for Spindly, CENP-C and DNA. Scalebar, 10 μm. (B) Scatter plot showing ratio of fluorescence intensity of Spindly to GFP-CENP-E_2055-2608_ wild type of mutants at spindle poles (WT n=70, E4A n=70, TS2AA n=74 and E4A TS2AA n=64). Bars represent median and 95% confidence interval. KS test with P ***<0.0001. (C) Representative immunofluorescence images of HeLa cells transfected with GFP-CENP-E_2055-2608_ wild type and mutants, stained for Mad1, CENP-C and DNA. Scalebar, 10 μm. (D) Scatter plot showing ratio of fluorescence intensity of Mad1 to GFP-CENP-E_2055-2608_ wild type of mutants at spindle poles (WT n=70, E4A n=62, TS2AA n=66 and E4A TS2AA n=670). Bars represent median and 95% confidence interval. KS test with P ***<0.0001. (E) Representative immunofluorescence images of HeLa cells transfected with GFP-CENP-E_2055-2608_ wild type and mutants, stained for Spindly, CENP-C and DNA. Scalebar, 10 μm. (F) Bar graph showing percentage of misaligned chromosomes in the presence of GFP-CENP-E_2055-2608_ (n=61), GFP-CENP-E_2055-2608_ E4A (n=52), GFP-CENP-E_2055-2608_ TS2AA (n=63) and GFP-CENP-E_2055-2608_ E4A TS2AA (n=56), data from 5 experiments. Mean and standard deviation are presented. Asterisks indicate ordinary one-way ANOVA test significance value. ****P<0.0001. (G) Schematic model explaining the two modes of recruitment of CENP-E to kinetochores in early mitosis and CENP-E removal in metaphase.

GFP-CENP-E_2055-2608_ has a dominant negative phenotype, causing chromosome misalignment, with over 80% of cells expressing the domain displaying misaligned chromosomes (Fig. 4E-F) (Chan et al., 1998; Legal et al., 2020). Around 40% of cells expressing either GFP-CENP-E_2055-2608_ E4A or GFP-CENP-E_2055-2608_ TS2AA mutants displayed misaligned chromosomes. In the presence of GFP-CENP-E_2055-2608_ E4A TS2AA, most chromosomes remained aligned, indicating the recruitment of corona-localized proteins to CENP-E_2055-2608_ prevents kinetochore alignment and formation of stable kinetochore-microtubule attachments, rather than the presence of CENP-E_2055-2608_ itself.

Our work reveals that CENP-E is recruited to unattached kinetochores through 2 separate pathways: a BubR1-dependent pathway and a pathway dependent on Dynein and the outer corona (Fig. 4G). The two domains involved in the respective interactions are close but structurally distinct and facilitate cooperative recruitment of CENP-E with assembly of the outer corona. We previously showed that the BubR1 recruitment pathway was kinetically fast and took place within minutes, while the second pathway was slower. It is likely that assembly of the outer corona, facilitated by Mps1 phosphorylation, needs to take place before CENP-E can bind to its receptor. As CENP-E is able to recruit the RZZ complex, Spindly and Mad1, we propose the BubR1-dependent pathway facilitates and could catalyze the assembly of the outer corona through CENP-E. Future studies are required to identify the molecular basis for CENP-E recruitment to the outer corona and how the force produced by the motor activity of CENP-E is transmitted to kinetochores to align them.

## Acknowledgements

We thank Jingchao Wu, Geert Kops, Susana Eibes, Marin Barisic, Verena Cmentowski and Andrea Musacchio for data and information sharing before publications. J. W. and O.D are supported by a Wellcome Senior Research Fellowship (207430 and 219413 respectively). JW is also a EMBO Young Investigator. We thank Dave Kelly for support in the COIL imaging facility and Rennos Fragkoudis and Peter Vegh at the Genome foundry. The Wellcome Trust Centre for Cell Biology is supported by core funding from the Wellcome Trust (203149).

## Methods

### Cloning

To assay the localisation in cell culture of CENP-E subdomains, the constructs were generated from CENP-E transcript variant 1 (NM_001813.2) and cloned into pBABE-puro containing an N-terminal GFP tag and using restriction enzymes (Cheeseman and Desai, 2005). Mutagenesis was performed using Quickchange site-directed mutagenesis kit (200523, Agilent) according to the manufacturer’s guidance. The full-length CENP-E constructs in pcDNA5 FRT/TO were assembled with the assistance of the Edinburgh Genome Foundry, a synthetic biology research facility specialising in the assembly of large DNA fragments at the University of Edinburgh. DNA fragments were synthesized by Geneart.

### Cell culture and experiments

Stable clonal HeLa cells lines expressing mCherry-CENP-A were generated as described previously using a retroviral system (Cheeseman and Desai, 2005). HeLa cells expressing mCherry-CENP-A were grown in DMEM (Life Technologies) supplemented with 10% FBS and Penicillin-Streptomycin (Gibco) at 37°C and 5% CO2. Cells were monthly checked for mycoplasma contamination (MycoAlert detection kit; Lonza). For live imaging, cells were plated on 35-mm glass-bottom microwell dishes (Ibidi). Transient transfections were conducted using Effectene reagent (QIAGEN) according to the manufacturer’s guidelines. Cells were imaged/fixed 24 to 48 hours later.

Knockdown of BubR1 and Dynein heavy chain (DHC) was achieved by transfecting 150 pM siRNA using Lipofectamine RNAiMax (Thermo fisher) according to the manufacturer’s instructions. Cells were fixed 72 hours later. The siRNA sequences used were: 5’-AAGGAUCAAACAUGACGGAAUUU-3’ for DHC and 5′-GCAATCAAGTCTCACAGAT-3’ for BubR1 (Espert et al., 2014). Treatment with GSK923295 was performed for 30 minutes at a concentration of 200 nM (Selleckchem). To generate detachable kinetochore crescents, cells were treated with 3.3 μM nocodazole (Calbiochem) for 3 hours followed by addition of 10 μM RO-3306 (Bio-Techne Ltd) for 25 minutes before fixing (Pereira et al, 2018; Sacristan et al, 2018).

### Cell lines

The Flp-In cell line expressing GFP-CENP-E_2055-2608_ was generated using HeLa T-REx Flp-In 293 cells according to the Flp-In system protocol (Thermo Fisher). GFP-CENP-E_2055-2608_ was cloned into pcDNA5 FRT/TO vector. After selection, the cell line was maintained in DMEM supplemented with tetracycline-free FBS (Gibco). Expression of GFP-CENP-E_2055-2608_ was induced with 1 μg/mL doxycycline for 24-36 hours (Sigma Aldrich).

### Immunofluorescence and microscopy

Cells were fixed with 3.8% formaldehyde in PHEM buffer (60 mM Pipes, 25 mM HEPES, 10 mM EGTA, 2 mM MgSO4, pH 7.0) or ice cold methanol for 20 minutes and stained using rabbit anti-spindly (1:200, Bethyl lab A301-354A), anti-Mad1 clone BB3-8 (1:1000, Merck), anti-ZW10 (Rabbit, Abcam ab21582), guinea pig anti-CENP-C (1:1000, MBL PD030) and mouse CENP-E (1:200, Abcam ab5093) antibodies. Hoechst 33342 (ThermoFisher Scientific; H3570) was used to stain DNA.

Images were recorded using a Deltavision core microscope (Applied Precision) equipped with a CoolSnap HQ2 CCD camera or widefield Eclipse Ti2 (Nikon) microscope equipped with a Prime 95B Scientific CMOS camera (Photometrics), using a 100x objective (CFI Plan Apochromat Lambda, 1.49 N.A). For fixed samples, Z-sections were acquired at 0.2-μm step size using 100X objective lens. For live-cell imaging, 10 Z-sections were acquired at 0.5-μm step size using 100X objective lens. Cells were imaged in Leibowitz L15 medium (ThermoFisher Scientific). Images were stored and visualized using an OMERO.insight client (OME) (Allan et al., 2012). Mean kinetochore fluorescence intensity within a circular ROI with an 8-pixel diameter was measured for with a background intensity recorded in an adjacent cytoplasmic area. Relative values for each kinetochore are calculated by subtracting the background values and dividing them by the background corrected CENP-C signal for that kinetochore. For quantification of pole accumulation of CENP-E, a similar approach was used. Mean pole fluorescence intensity, marked by the bipolar CENP-E_2055-2608_ localization within a circular ROI was measured for with a background intensity recorded in an adjacent cytoplasmic area. Average intensity was calculated by subtracting the background values and dividing them by the background corrected pole-localized CENP-E signal for that kinetochore. For ZW10, Mad1 and Spindly intensity, the ROI from the CENP-E channel was used, and average intensity was calculated as above. Data were analysed using ImageJ (Schneider et al., 2012). Images for immunofluorescence were deconvolved using softWoRx (Applied Precision).

### Protein expression and crystallization

His6-CENP-E_2452-2598_ was cloned into pET3aTr and expressed in BL21 cells and purified as previously described (Legal et al., 2020). Protein was concentrated to 11 mg/ml and crystallized in 0.2M Potassium citrate tribasic monohydrate, 20% w/v polyethylene glycol 3350 pH8.3 within a few days at room temperature. The crystal was protected in 30% glycerol mixed with mother liquor and frozen in liquid nitrogen.

### Data collection and structure determination

X-ray diffraction data were collected at 0.9795 Å, 100 K, as 3600 consecutive 0.10° frames of 0.050 s exposure on a Dectris Eiger2 XE 16M detector at beamline I04 of the Diamond Light Source synchrotron facility (Oxfordshire, UK). Data were indexed, integrated in XDS (Kabsch, 2010), scaled in Aimless (Evans, 2011), and merged with anisotropic correction and ellipsoidal truncation by STARANISO (Tickle, 2018), using AutoPROC (Vonrhein et al., 2011). Crystals belong to tetragonal spacegroup P4_2_2_1_2 (cell dimensions a = 84.02 Å, b = 84.02 Å, c = 65.02 Å, α = 90°, β = 90°, γ = 90°), with a CENP-E dimer in the asymmetric unit. Structure solution was achieved through fragment-based molecular replacement using ARCIMBOLDO_LITE (Rodriguez et al., 2009), in which six helices of 14 amino-acids were placed by PHASER (McCoy et al., 2007) and extended by tracing in SHELXE utilising its coiled-coil mode (Caballero et al., 2018). A correct solution was identified by a SHELXE correlation coefficient of 56.4%. Model building was performed through iterative re-building by PHENIX Autobuild (Adams et al., 2010) and manual building in Coot (Emsley et al., 2010). The structure was refined using PHENIX refine (Adams et al., 2010), using isotropic atomic displacement parameters with one TLS group per chain. The structure was refined against data to anisotropy-corrected data with resolution limits between 2.14 Å and 2.71 Å, to R and R_free_ values of 0.2161 and 0.2455 respectively, with 100% of residues within the favoured regions of the Ramachandran plot (0 outliers), clashscore of 1.05 and overall MolProbity score of 0.81 (Chen et al., 2010). PDB entry is: 8OWI.

### Structural modelling

Models were generated using a local installation of *Alphafold2* (Jumper et al., 2021) (Evans et al., 2021). The BubR1-targeting domain (amino-acids 2055-2377) was modelled as a monomer using the monomer pipeline. The corona-targeting domain (amino-acids 2356-2607) was modelled as a dimer using the multimer pipeline, with the newly solved crystal structure (PDB accession 8OWI) specified as a template. The resultant structures were aligned based on their overlapping sequences, and were joined to form two full chains, and their intervening linkers were fitted using geometry and Ramachandran restraints. *Alphafold2* multimer modelling data were analysed using modules from the ColabFold notebook (Mirdita et al., 2022). Models were edited, combined and flexible linkers were remodelled using the *PyMOL* Molecular Graphics System, Version 2.0.4 Schrödinger, LLC, and *Coot* (Emsley et al., 2010).

### Statistics and reproducibility

Statistical analyses were performed using Prism 9 (GraphPad Software). For the calculation of the error on the median, we report the upper 95% confidence interval. No statistical method was used to predetermine sample size. No samples were excluded from the analyses. The investigators were not blinded to allocation during experiments and outcome assessment. All experiments were performed and quantified from 2-3 independent experiments and representative data are shown.

### Western blots

All western blots performed used the antibodies listed in the table below at the indicated concentrations.

| Antibody | Source | Reference | Concentration WB |
| --- | --- | --- | --- |
| Tubulin | Mouse | Sigma, T8328 | 1:1000 |
| BubR1 | Rabbit | Bethyl Lab, A300-386A | 1:2000 |
| DHC | Rabbit | Atlas antibodies, hpa003742 | 1:1000 |

---

**Supplementary figure 1:**
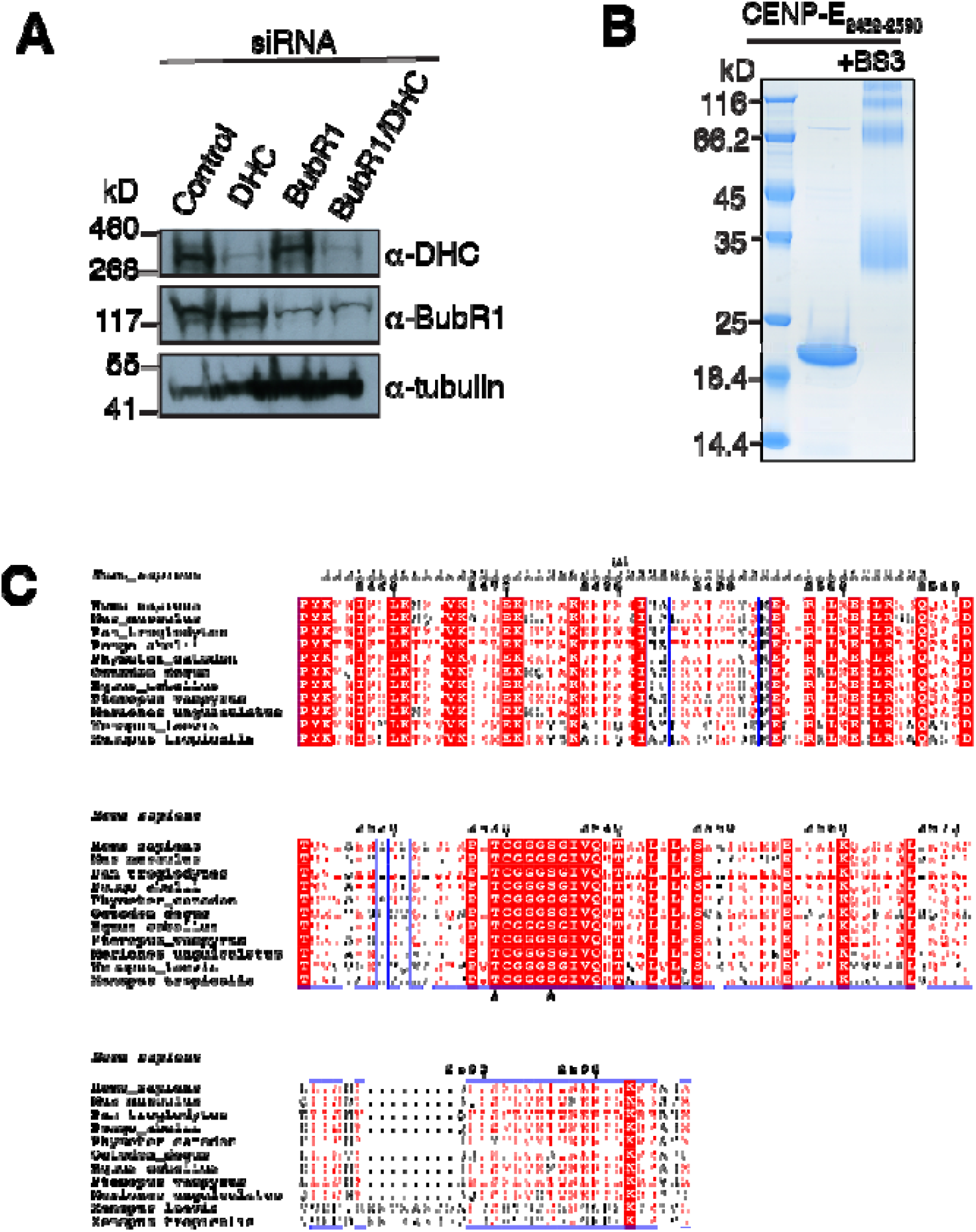
(A) Western blot for cells in Figure 1 after siRNA treatment probed for BubR1/DHC and tubulin as a loading control. (B) Coomassie-stained gel showing CENP-E_2452-2598_ and after crosslinking with BS3. (C) Sequence alignment of the human CENP-E_2287-2246_ with mouse, chimpanzee, orangutan, sperm whale, degu, horse, flying fox, gerbil and Xenopus. Boxed red and blue are the conserved and similar amino acids across all species, respectively. Amino acids in red are those with conserved properties in at least 3 sequences. The glutamates necessary for outer corona binding are marked with a star (*).

**Supplementary figure 2:**
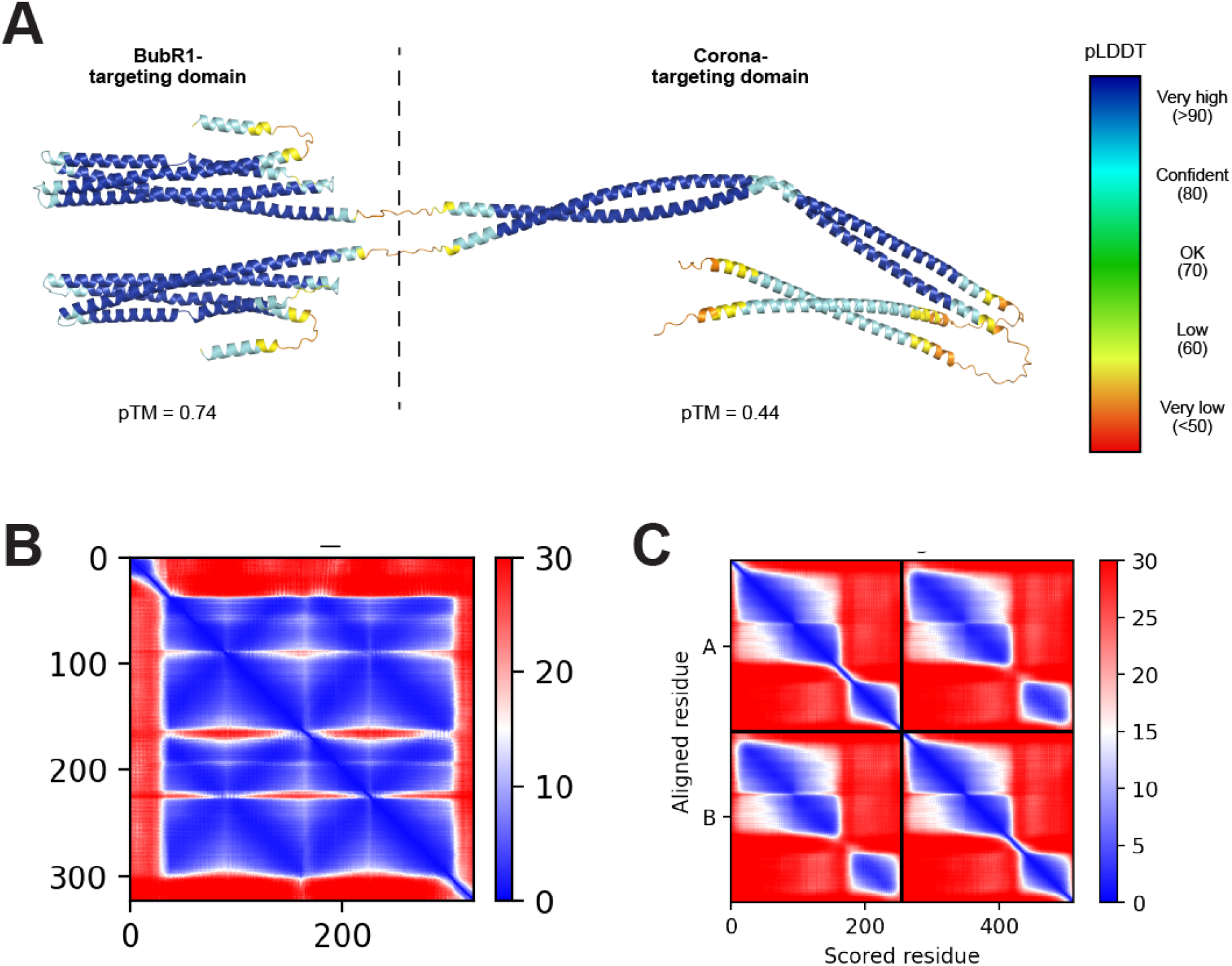
(A) Model of the CENP-E_2055-2608_ coloured according to pLDDT scores, between blue (>90) and red (<50). (B) Predicted aligned error scores between each amino-acid of the single chain and two chains of the BubR1- and corona-targeting domains of CENP-E, respectively, between blue (low error) and red (high error).

**Table 1:** table summarizing crystallographic data for the structure of CENP-E_2454-2494_.

**Table 2:** table summarizing the plasmids used in this study.

